# Cross-generational plasticity in Atlantic silversides (*Menidia menidia*) under the combined effects of hypoxia and acidification

**DOI:** 10.1101/2024.05.22.595394

**Authors:** Christopher S. Murray, Ayanna Mays, Matthew Long, Neelakanteswar Aluru

## Abstract

We investigated the potential for cross-generational plasticity to influence how offspring respond to hypoxia and ocean acidification (hereafter HypOA) in the coastal forage fish Atlantic silverside (*Menidia menidia*). Mature wild silversides were treated with a control [dissolved oxygen (DO):100% air saturation (a.s.) / *p*CO_2_: 650 µatm] or HypOA conditions [DO: 40% a.s. / *p*CO_2_: 2300 µatm] for 10 days prior to spawning. Their offspring were reared under both treatments in factorial experimental design. Parental acclimation to HypOA altered several offspring traits, including increased embryo survival under HypOA and an overall reduction in post-hatch growth rate. Offspring from HypOA-treated parents that were reared under control conditions had larger eyes across the developmental period. When compared against the overall control group, larvae directly exposed to HypOA exhibited 2,416 differentially expressed transcripts (DETs). Although most of these DETs were specific to individual parental treatments, the most enriched Gene Ontology terms were conserved across parental treatments, including terms related to neurotransmitter secretion, nervous system development, axon pathfinding, calcium channel activity, proteolysis, and extracellular matrix organization. Larvae from HypOA-treated parents that were reared under control conditions exhibited a shift in constitutive gene expression similar to that seen in larvae directly exposed to HypOA. This highly consistent finding indicates that parental acclimation before fertilization promotes the transcriptional frontloading of genes in offspring. This effect may have primed regulatory functions in offspring that sense and respond to low DO and elevated *p*CO_2_ conditions. Though, our results suggest that this altered developmental phenotype may have some negative fitness consequences for offspring.

## 1 Introduction

Hypoxia and acidification (hereafter HypOA) are naturally co-occurring abiotic stressors in productive coastal marine habitats, driven by shared physical and biochemical processes (Rabalais et al., 2009; Wallace et al., 2021). Human-induced factors like eutrophication and habitat degradation have increased the severity, duration, and spatial scale of HypOA in many coastal areas, and global warming and ocean acidification will exacerbate this trend (Cai et al., 2021; Keeling et al., 2010; Long and Mora, 2023). Nearshore marine ecosystems are important spawning and nursery habitat for estuarine and marine fish, thus the intensification of these stressors directly threatens the viability of many ecologically and economically important populations (Baumann, 2019; Breitburg, 2002). Exposure to low dissolved oxygen (DO) has been shown to adversely impact reproductive and developmental processes (Richards, 2009) and many fish species show sublethal effects at concentrations well above typical operational hypoxia thresholds of 2 - 3 mg L^-1^ (Wu, 2009). Experiments that have measured hypoxia tolerance have largely overlooked the co-occurrence of hypoxia and acidification (Breitburg et al., 2019; Gobler and Baumann, 2016). Recent experiments on fish have shown additive and complex interactive effects resulting from combined low DO and elevated *p*CO_2_, highlighting the importance of multi-stressor studies to correctly assess organismal sensitives (Cross et al., 2019; DePasquale et al., 2015; Miller et al., 2016; Schwemmer et al., 2020).

Cross-generational plasticity could play an important role in how fish populations respond to worsening HypOA. It describes how the environmental experiences of a parent can influence the phenotype of their offspring through various “non-genetic” factors of inheritance (Adrian-Kalchhauser et al., 2020; Byrne et al., 2020). In fish, these factors include classically described maternal effects, such as the provisioning of eggs with nutrients, hormones, proteins, mRNA, and other cytoplasmic components that can influence offspring development (Green, 2008). Inheritable epigenetic factors, including DNA methylation, histone modification, and small noncoding RNAs, can be transmitted through both maternal and paternal germlines and influence offspring gene expression and responses to environmental stress (Ben Maamar et al., 2018; Kelley et al., 2021; Ord et al., 2020). Cross-generational plasticity likely evolved as a fitness-enhancing strategy in organisms that reproduce in variable environments, enabling parents to anticipate conditions offspring are likely to encounter and provide a developmental advantage (Bonduriansky, 2021). For example, treating adult zebrafish to hypoxia prior to spawning increases the acute hypoxia tolerance of offspring (Ho and Burggren, 2012; Ragsdale et al., 2022). Such a capability would be advantageous in the shallow, slow-moving waters of natural zebrafish habitat, where diurnal hypoxia is common (Spence et al., 2008). Similarly, damselfishes experience natural *p*CO_2_ fluctuations in coral reef habitats, and beneficial cross-generational effects to ocean acidification have been observed in multiple species (Miller et al., 2012; Monroe et al., 2021).

Transcriptional frontloading may be a useful framework for conceptualizing how parental effects alter gene expression patterns in offspring. It defines a type of acclimatory gene expression that occurs when exposure to priming stressor elicits a persistent shift in the expression of stress response genes, enabling the organism to initiate a more rapid and efficient adaptive response upon subsequent challenges within its lifetime (Barshis et al., 2013; Hackerott et al., 2021). For example, reef-building corals (*Acropora hyacinthus*) from thermally variable environments show a frontloading of genes involved in thermal stress and exhibit a greater tolerance to bleaching compared to conspecifics from deeper, more thermally stable environments (Barshis et al., 2013). Likewise, Juvenile Pacific geoduck clams (*Panopea Generosa*) that were exposed to elevated *p*CO_2_ during early development showed a frontloading of genes involved in maintain homeostatic functions under acidification and consequently were more tolerant of repeated exposures later in life (Gurr et al., 2022). However, this framework has not yet been widely applied in detecting transcriptional effects associated with cross-generational plasticity.

Atlantic silverside (*Menidia menidia*) is a widespread and highly abundant forage fish that inhabits coastal ecosystems along the North American eastern seaboard (Hice et al., 2012; Middaugh et al., 1987). Silversides are an annual asynchronous batch-spawning species that reproduce multiple times at two-week intervals during spring and summer. Thus, offspring from successive spawning events encounter very different environmental conditions as nearshore systems become more susceptible to HypOA due to rising temperatures and increased biological productivity (Baumann et al., 2015). Although hypoxia is a natural component of their developmental environment, the early life stages of Atlantic silversides are relatively sensitive to low DO compared to other fish species with similar habitat preference (DePasquale et al., 2015; Dixon et al., 2017). Furthermore, simultaneous exposure to elevated *p*CO_2_ can increase hypoxia sensitivity (Cross et al., 2019; DePasquale et al., 2015; Miller et al., 2016; Schwemmer et al., 2020). Intriguingly, the tolerance of wild silverside offspring to elevated *p*CO_2_ varies seasonally, with subsequent cohorts exhibiting an increasing CO_2_ tolerance that coincides with seasonally progressing coastal acidification (Baumann et al., 2018; Murray et al., 2014). The underlying mechanism for this effect remains uncertain—whether it arises directly from the exposure of spawning adults to increasing daily average or maximum *p*CO_2_, or if it is linked to other seasonal changes in parental and offspring phenotype. Furthermore, the potential impact of cross-generational plasticity on offspring hypoxia tolerance in this species has yet to be explored.

The objective of this study was to investigate the potential for cross-generational plasticity to influence the responses of Atlantic silverside offspring to HypOA. Wild reproductively mature Atlantic silversides were treated for 10 days to control conditions or a combined HypOA treatment. Their offspring were reared in a factorial experimental design with four offspring treatment groups: control-treated parents and control-reared offspring (CC), control-treated parents and HypOA-reared offspring (CH), HypOA-treated parents and control-reared offspring (HC), and HypOA-treated parents and HypOA-reared offspring (HH) (Fig. 1). We monitored survival and growth during the first 19 days of embryo and larval development, by which time notochord flexion is typically completed. At trial termination, surviving larvae were sampled for RNA sequencing, to characterize how parental treatment to HypOA influenced larval gene expression. We hypothesized that offspring fertilized by HypOA-treated parents will show increased survival and growth under the combined stressor. Furthermore, we hypothesized that responses in offspring would be underpinned by a frontloading of distinct genes involved in functional pathways that promote tolerance to low DO and elevated *p*CO_2_.

**Fig. 1.**
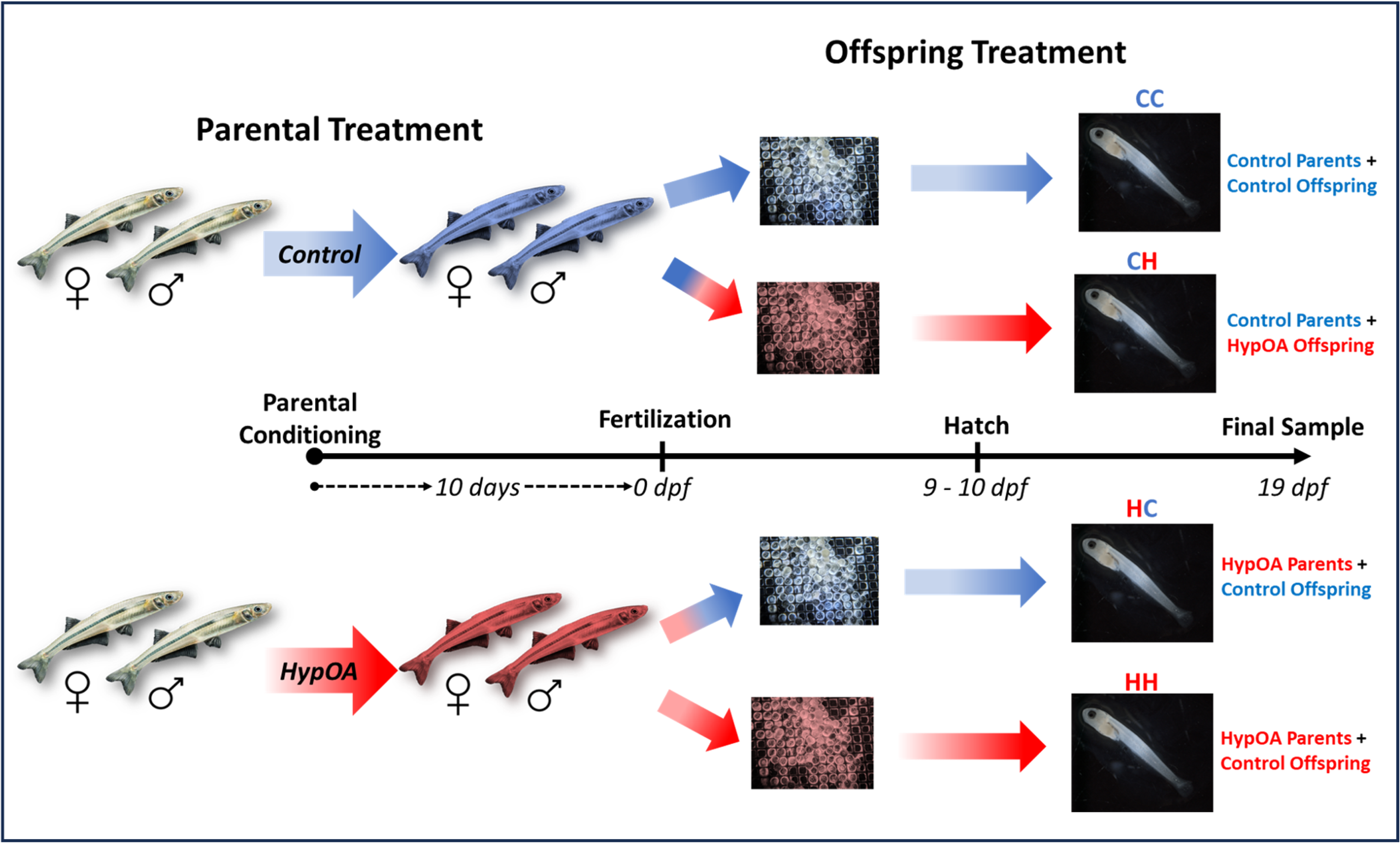
Schematic of the experimental design. Wild mature Atlantic silversides were treated to control or a combined hypoxia and acidification (HypOA) treatment for 10 days. After fertilization, their embryos were split and reared under both offspring treatments in a factorial design to create four offspring treatment groups. Larvae were counted and sampled at hatch (9 – 10 dpf) and surviving larvae were recounted and sampled at 19 dpf to measure post-hatch survival and growth responses. RNA was extracted from an additional set of larvae at 19 dpf for transcriptomic analysis.

## 2 Methods

### 2.1 Experimental treatments

We implemented two experimental treatment levels: a control treatment with a DO set to 100% air saturation (a.s.) (8.9 mg L^-1^) and 650 µatm *p*CO_2_ (7.83 pH_Total scale_), and a static HypOA treatment with a DO at 40% a.s. (3.6 mg L^-1^) combined with 2,300 µatm *p*CO_2_ (7.33 pH_T_). The control treatment reflects the typical daytime DO and *p*CO_2_ conditions observed in a Atlantic silverside spawning habitat during their reproductive season (Baumann et al., 2015). Conversely, the HypOA treatment reflects conditions that occur infrequently in nearshore temperate environment during periods of extreme net heterotrophy (Baumann and Smith, 2017; Baumann et al., 2015; Wallace et al., 2021), that will likely become more common in future (Breitburg et al., 2019; Cai et al., 2021). This combination of low DO and elevated *p*CO_2_ has been shown to reduce the survival and developmental rates of silverside embryos and larvae (Cross et al., 2019).

### 2.2 Description of the experimental system

The experiment was conducted at the Environmental Systems Laboratory (ESL) facility at the WHOI in an automated experimental seawater system comprised of eight 900-L circular tanks (four tanks each for control and HypOA treatment) and one 600-L mixing tank. A schematic of the experimental setup is shown in Fig. S1. Each 900-L tank was outfitted with multiple 20-L rearing containers (white buckets fitted with six 3-cm flowthrough holes covered in 300-µm screening) to accommodate embryos and larvae. The four control tanks received temperature-controlled seawater (21°C) sourced from Vineyard Sound (sand-filtered, UV-sterilized, and maintained at the control treatment DO and *p*CO_2_). HypOA treatment conditions were maintained within the mixing tank using custom software developed in Arduino IDE and operated by a Teensy 4.1 Development Board. Briefly, the mixing tank received a continuous flow of ambient seawater, with DO, pH, and temperature measured every 10 seconds by Pyroscience Ultra Compact (PICO) DO and pH meters, fitted with fiberoptic DO or pH sensors and PT100 temperature sensors. Calibration of the probes was performed weekly according to the manufacturer’s guidelines using oxygen and pH_T_ calibration solutions provided by Pyroscience. HypOA levels were regulated through a simple feedback mechanism: when the current DO or pH exceeded the treatment set point, the software opened a solenoid valve to introduce compressed gas (N_2_ or CO_2_) into ultrafine gas-diffusers positioned at the bottom of the mixing tank until the setpoint was reached. The software was optimized to implement brief and precisely controlled gas additions, ensuring stable DO and pH conditions in the mixing tank. Adjusted HypOA seawater was continuously pumped to the main tanks and individual rearing containers.

DO levels in individual rearing units were cross verified using a handheld Pyroscience Firesting-GO2 meter fitted with an optical DO sensor. A two-point calibration (100% and 0% DO) was performed daily using air-saturated seawater and a concentrated NaSO_3_ seawater solution. pH conditions were verified via a Hach HQ11D pH meter fitted with a Hach PHC28101 IntelliCal pH probe. The probe was calibrated daily using pH_T_ buffers from PyroScience. To validate target *p*CO_2_ conditions, seawater was sampled every 1 – 2 days from one of the replicate rearing containers per offspring treatment group, resulting in 12 seawater samples per offspring treatment group during the 19-day experiment. Seawater samples were sterilized with mercuric chloride and stored in sealed bottles for later analysis of carbon chemistry parameters. Temperature and salinity (via refractometer) were recorded at the time of sampling. Samples were measured in triplicate for dissolved inorganic carbon (DIC) using an Apollo AS-D1 analyzer connected to a Picarro G-2121i cavity ringdown system. Total alkalinity (*A*_T_) was measured using an open-system Gran titration on 5-ml samples in triplicate, using a Metrohm 805 Dosimat and a robotic Titrosampler. Both systems were calibrated against seawater Certified Reference Materials (Dickson, 2024). The remaining *in situ* carbon chemistry parameters of pH_T,_ *p*CO_2_, bicarbonate, and carbonate ions were calculated using the R package seacarb (Gattuso et al., 2020) based on direct measurements of DIC, *A*_T_, and the salinity and temperature at the time of water sampling. Equilibrium constants for the dissociation of carbonic acid in seawater (K1 and K2) followed Mehrbach et al. (1973) and refitted by Dickson and Millero (1987). The constant for KHSO_4_ followed Dickson (1990). Measured and derived carbon chemistry parameters are listed in Table 1.

**Table 1:**
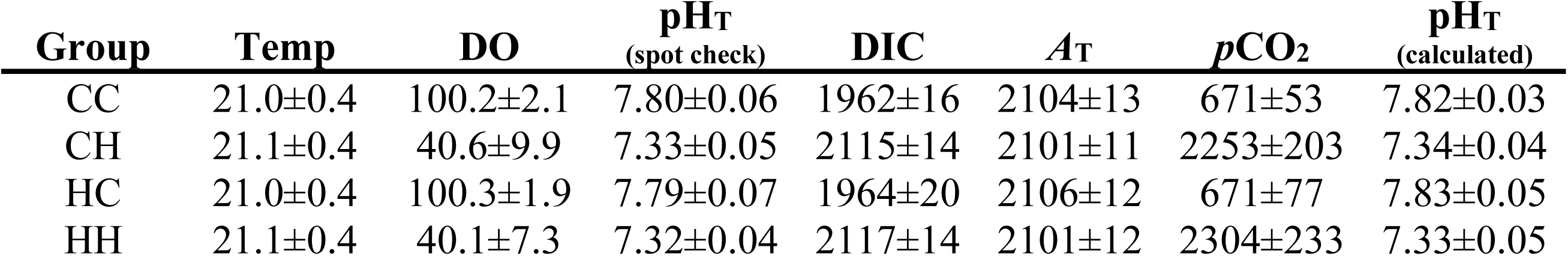
Summary of treatment conditions. . Values are presented as offspring treatment means (± s.d.). Temperature (temp, °C), dissolved oxygen (DO, % air saturation), and pH_T (spot check)_ from daily spot check measurements using handheld dip probes (N = 20). Total alkalinity (*A*_T_, μmol kg^-1^) and dissolved inorganic carbon (DIC, μmol kg^-1^) were measured directly from preserved seawater samples (N = 12). *p*CO_2_ (μatm) and pH_T (calculated)_ were calculated from measured DIC, *A*_T_, temperature, and salinity.

### 2.3 Wild fish collection and care

Animal welfare protocols and experimental procedures followed guidelines set forth by WHOI’s Institutional Animal Care and Use Committee (WHOI protocol #28188). Wild adult Atlantic silversides were collected on May 31, 2022, by beach seine (30 × 2 m) from a single location in Mumford Cove, CT, a small seagrass dominated embayment connected to eastern Long Island Sound (41.32°N, 72.02°W). Fish were transported in an aerated cooler to the ESL where they were separated by sex and then into two groups for treatment to either control or HypOA conditions (34 males and 37 females per treatment group). Adults were acclimated to laboratory conditions for five days in large circular tanks (one tank per sex and treatment group). Tanks received a continuous flow (5 L min^-1^) of ambient seawater. Fish were hand fed to satiation twice daily with frozen adult brine shrimp and blood worms (San Francisco Bay Brand).

### 2.4 Parental acclimation and fertilization protocols

Following the initial five-day acclimation period, adult fish were exposed to their respective experimental treatment conditions for 10 days. Parental treatment to HypOA involved exposure to a fluctuating DO and *p*CO_2_ regime [control treatment: 7 AM – 7 PM; HypOA treatment: 7PM – 7AM], mimicking the diurnal fluctuations and nocturnal hypoxia and acidification that characterize nearshore marine environments (Baumann et al., 2015). Continuous DO measurements of adult HypOA tanks and DO and pH conditions in the mixing tank are displayed in Figure S2. We implemented fluctuating HypOA protocol over a static treatment out of concerns that chronic exposure to HypOA conditions might strongly affect reproductive success. Adults were hand fed frozen brine shrimp and blood worms to satiation twice daily. Following acclimation, separate *in vitro* fertilization (IVF) were conducted for each parental group, following established protocols (Murray and Baumann, 2018; Murray et al., 2017). These methods are optimized to produce maximum genetic diversity within experimental replicates (Malvezzi et al., 2015). For each parental group, 17 - 20 males and 7 females contributed gametes during fertilization. Their lengths and weights are reported in Table S1. Embryos were screened within an hour after IVF and viable embryos were selected for rearing.

### 2.5 Offspring rearing and sampling

Embryos were reared in 500-ml plastic cups with 300-µm screened bottoms that were floated inside the 20-L rearing containers. For calculating survival, 50 embryos were used per replicate (N = 4 replicate containers per offspring treatment group). For measuring growth and transcriptomic analysis, 200 embryos were used (N = 4 replicate containers per offspring treatment group) (Fig. S1). Seawater lines were positioned directly inside the embryo cups to provide a direct seawater flow over embryos (30 ml min^-1^). Initially, all rearing containers received a flowthrough of control seawater and the introduction of HypOA-adjusted seawater into designated replicates began four hours post fertilization. Spot-check measurement of all rearing containers for DO, pH, and temperature were made at least once daily using handheld GO_2_ and Hach pH meters. A summary of the spot-check measurements is provided in Table 1.

Starting 6 days post-fertilization (dpf), replicates were checked in the morning for newly hatched larvae. Hatchlings from survival replicates were counted and moved to a new 20-L container to monitor post-hatch survival (flowthrough increased to 120 ml min^-1^). Subsamples for morphometric analysis at hatch (N = 10) were collected on the first morning when a growth replicate had at least 20 new hatchlings. The larvae were euthanized by an overdose of MS-222 and were fixed in 4% neutrally buffered formalin for 24 h and transferred to 75% ethanol for storage. The remaining hatchlings from growth replicates were moved to a new 20-L container for continued rearing. Larvae were fed daily with newly hatched brine shrimp nauplii (San Francisco Bay strain) at an initial concentration of 2-3 nauplii ml^-1^. Replicate vessels were siphoned daily for uneaten food and waste. The continuous seawater flow through ensured that ammonia levels remained near 0 ppm, verified daily via API Ammonia Test Kit. At 19 dpf (9 or 10 days post hatch (dph) based on the treatment and hatch timing) the survival replicates were recounted to determine the final survival rate. Larvae were subsampled for morphometric measurements (N = 12) as described above. Additionally, a subset of larvae (19dpf) was sampled individually (N = 6 per treatment, two larvae from three of the replicate rearing containers) and immediately snap-frozen in liquid nitrogen and stored at −80°C for subsequent RNAseq analysis.

Formalin-preserved larvae were laterally photographed using a digital camera attached to a dissecting microscope. Morphometric measurements were made using ImageJ2 (V. 1.53; Rueden et al., 2017). Newly hatched larvae were measured for four traits: standard length (tip of the snout to the end of the notochord; nearest 0.01 mm), eye diameter (anterior to posterior diameter of left eye, 0.01 mm), body depth (myomere height immediately posterior to the anus, 0.01 mm), and yolk sac profile area (0.01 mm^2^). The larvae sampled at 19 dpf were measured for standard length, body depth, and eye diameter.

### 2.6 RNA extraction and sequencing

Total RNA was extracted from individual larvae (N = 6 per offspring treatment group; 4 treatments) using the Qiagen RNeasy Mini extraction kit using the manufacturers guidelines. RNA purity was confirmed using a Nanodrop 2000 Spectrophotometer and total RNA concentration was quantified using a Qubit fluorometer. The RNA samples were shipped on dry ice to Novogene Inc. (Sacramento, CA), where quality was assessed using Agilent Bioanalyzer (RIN > 9.3 for all samples). NEBNext Ultra™ II Directional RNA Library Prep Kit was used for library generation and sequenced on a NovaSeq 6000 platform (150 bp paired-end reads) at a sequencing depth of 20 million reads per sample.

### 2.7 Bioinformatics

Initial data processing involved trimming reads to remove adapter sequences, reads with unknown bases, and low-quality reads using the default settings for paired-end data in Fastp v0.23.4 (Chen et al., 2018). Sequencing produced an average of 17.74±1.39 million clean reads per sample. The STAR v.2.6.1d tool (Dobin et al., 2013) was used to align reads to a high-quality *M. menidia* reference genome (Jacobs et al., 2022). Gene count quantification for each sample was performed using HTseq-count v.0.11.1 (Anders et al., 2015). On average, 73.0±1.2% of clean reads were mapped to the *M. menidia* genome across samples, resulting in the annotation of 17,263 unique coding sequences (transcripts). We focused our analysis on 15,253 transcripts with annotations for NCBI gene symbols and Gene Ontology terms (The Gene Ontology Consortium, 2018).

### 2.8 Statistical analysis of survival and growth traits

Statistical procedures were completed using R (v. 4.0.2) in RStudio (v 1.3). Model performance and adherence to assumptions were confirmed using the *Performance* package (Lüdecke et al., 2020). Statistical significance was set at α = 0.05. Results are presented as treatment means ± s.d. unless specified otherwise. Time (days) to peak hatch was calculated for each replicate (i.e., the day with the highest hatch count). Replicate survival was quantified for two intervals: embryo survival (ratio of total hatch count over the initial 50 embryos) and larval survival (ratio of larvae surviving at 19 dpf over initial hatch count). Ratio data were logit transformed prior to testing (Warton and Hui, 2011). The individual and interactive effects of offspring × parental treatment conditions on peak hatch and survival traits were examined using two-way analysis of variance (ANOVA).

The individual and interactive effects of offspring × parental treatments on standard length at hatch, yolk sac area at hatch, and standard length a trial termination were analyzed using mixed-effects ANOVAs (Satterthwaite’s degrees of freedom method) using the *lme4* and *lmerTest* R packages (Bates et al., 2015; Kuznetsova et al., 2017). Replicate ID was set as a random factor to account for the shared environment of larvae sampled from the same rearing container. We calculated size-independent indices for body depth and eye width. Standardized residuals were derived from a common linear regression fit with standard length using all samples pooled across treatments for each age group (Fig. S3). We used mixed-effect ANOVAs, as described above, to test treatment effects on normalized body depth and normalized eye width for larvae of each age. A larval growth rate (mm × d^-1^) was computed for each subsample replicate as the average final standard length minus the average hatch standard length divided by the growth interval in days. Treatment effects on larval growth rates were analyzed by two-way ANOVA. Post-hoc multiple-comparison tests were conducted using Tukey’s HSD via the R package *emmeans* (Lenth, 2018).

### 2.9 Statistical analysis of RNAseq data

Aligned gene counts from all samples were assembled into a gene-level DGEList, filtered, and normalized using the R package *edgeR* v.3.40.2 (Robinson et al., 2010). Lowly expressed genes were filtered out using function filterByExpr (default parameters). Raw counts were normalized (calcNormFactors function, TMM method) and PERMANOVA (1*e*^6^ permutations) was used to test significant effects of parental and offspring HypOA exposure on global gene expression. Sample ordination was visualized using PCoA (Manhattan distances) in the *vegan* package (Oksanen J et al., 2023). Differentially expressed transcripts (DETs) were identified using genewise negative binomial generalized linear models (glmQLFit and glmQLFTest functions). Significance of DETs was determined at a false discovery rate (FDR) of < 0.05.

To determine the functional patterns associated with changes in gene expression, we assessed DETs from different sets of pairwise comparisons for significantly enriched (*p value* < 0.05) GO biological process (BP) and GO molecular function (MF) terms using the enricher function in *ClusterProfiler* v.4.6.2 (Wu et al., 2021). The background gene list included all expressed transcripts in the filtered DGEList. Redundancy reduction amongst enriched terms was carried out using the R package *rrvgo* (Sayols, 2023). A similarity matrix was generated based on the frequency of shared genes between GO terms using the calculateSimMatrix function (Wang method) and similar terms were clustered into groups using the reduceSimMatrix function (similarity threshold of 0.7). Parent terms for each cluster were chosen based on the term with the lowest *p value*. Heatmaps were plotted using the R package *complexHeatmap* (Gu et al., 2016).

To better understand how parental environmental influenced the transcriptional plasticity of offspring, we examined all unique DETs identified in the CHvs.CC and HHvs.CC pairwise comparisons (N = 2,416) for patterns of transcriptional frontloading using methods adapted from Barshis et al. (2013). To estimate how parental treatment to HypOA influenced the constitutive gene expression of offspring reared under control conditions, a ‘constitutive expression ratio’ was calculated as gene expression (normalized and variance-stabilized counts per million (CPM)) of HC larvae divided by the gene expression of CC larvae (CPM_HC_ / CPM_CC_). Next, we calculated the raw fold change values (FC) separately for upregulated (FC = 2^log2FC^) and downregulated DETs (FC = .5^log2FC^). FCs were used to calculate a ‘fold change ratio’ for each DETs (FC_HHvsHC_ / FC_CHvsCC_) to quantify how parental acclimation influenced gene expression plasticity between offspring treatments. Both ratios were log-transformed to improve visualization. Genes were classified into three response groups: (1) *Frontloaded* – transcripts identified as having a reduced response in larvae from HypOA-treated parents compared to larvae from control parents (fold change ratio < 0). These transcripts exhibited constitutive expression ratio > 0 for upregulated DETs or a constitutive expression ratio < 0 for downregulated DETs. (2) *Increased Plasticity* - transcripts with greater FC responses in larvae from HypOA-treated parents. (3) *Dampened* - transcripts with lower fold change responses between offspring treatments from HypOA-treated parents that coincided with a change in constitutive expression that opposed the effect of direct HypOA exposure (increased and decreased constitutive expression for down- and upregulated DETs, respectively). Two-way ANOVAs were used to test for treatment effects on the gene expression (normalized and variance-stabilized log_2_CPM) withing gene categories. GO enrichment analysis of frontloaded, increased plasticity, and damped transcripts was carried out using the methods described above.

## 3 Results

### 3.1 Survival and hatching

Embryo survival was significantly affected by a parental × offspring interactive effect (ANOVA, *p =* 0.017, Fig. 2A). HypOA exposure significantly reduced the survival of embryos from control parents (Tukey’s HSD, *p* = 0.006, Fig 2A). However, there was no difference in the survival rates of embryos reared under HypOA and control conditions from HypOA-treated adults (Tukey’s HSD, *p* = 0.9901, Fig. 2A). Time to peak hatch was 9 dpf in embryos reared under the control treatment and this was delayed by one day (10dpf) in embryos exposed to HypOA (ANOVA, *p* = 0.001). Parental treatment had no effect on time to hatch. Larval survival was generally high and not significantly affected by offspring treatment but was significantly lower in offspring from HypOA exposed parents when averaged across offspring treatments (ANOVA, *p* = 0.022, Fig. 2B).

**Fig. 2:**
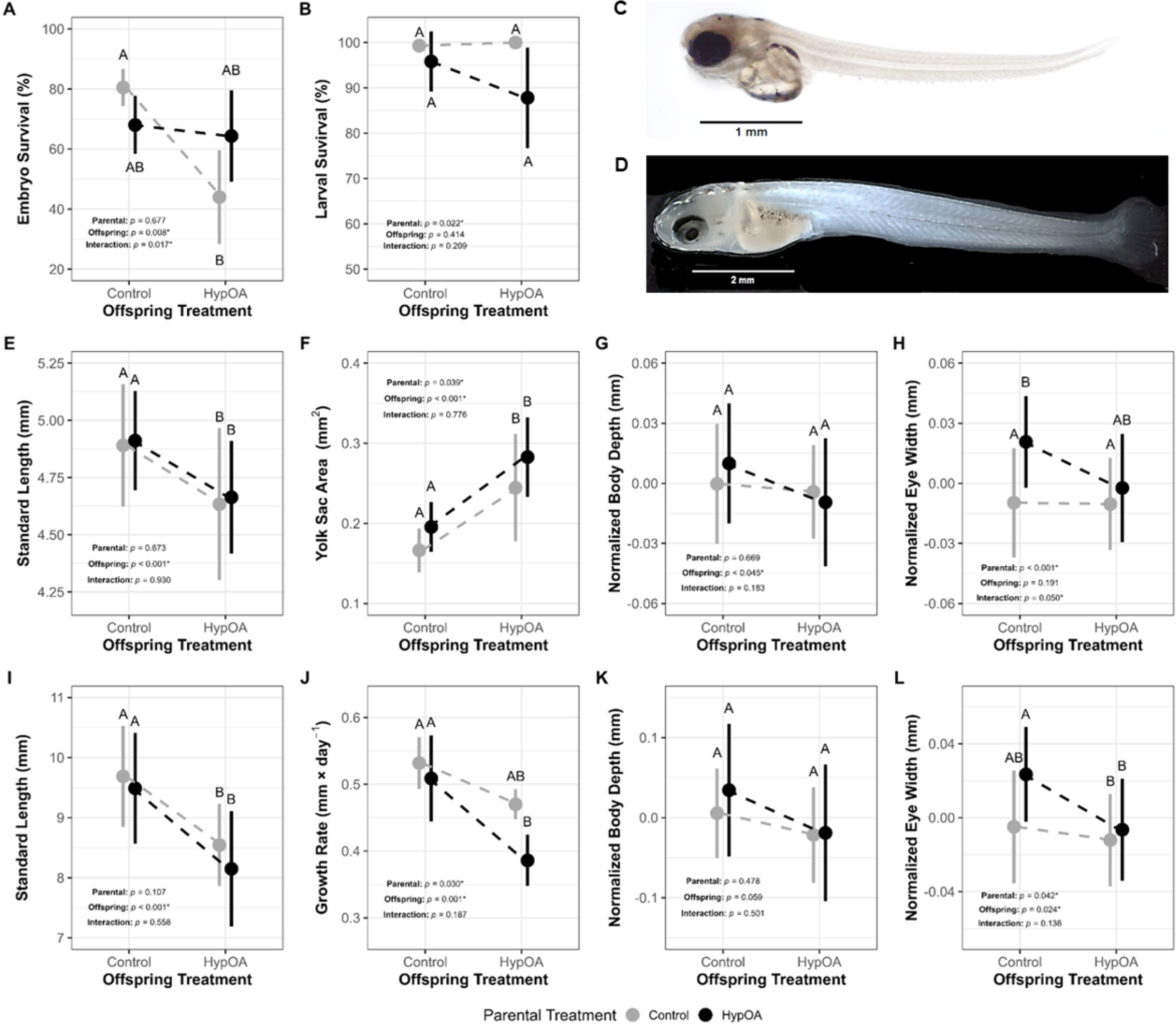
Treatment effects on offspring survival and morphometric traits. (A) Embryo survival (%, fertilization to hatch, N = 4) and (B) post-hatch larval survival (%, hatch to 19 days post-fertilization (dpf), N = 4). Images of (C) newly hatched larvae (9 dpf) and (D) larvae at 19 dpf or 10 days post-hatch. (E) Standard length at hatch (mm, N = 10); (F) yolk sac profile area at hatch (mm^2^, N = 10); (G) normalized body depth at hatch (mm, N = 10); (H) normalized eye width at hatch (mm, N = 10); (I) final standard length (mm, N =12); (J) post-hatch growth rate (mm d^-1^, N = 4); (K) final normalized body depth (mm, N = 12); (L) final normalized eye width (mm, N = 12). Large circles show treatment mean values and vertical lines indicate ± s.d. calculated from treatment mean values. Hashed lines show reaction norms by parental treatment. Significance levels (*p* values) are presented from two-way ANOVAs (survival and growth rate) or two-way mixed effect ANOVAs (morphometric traits) testing the effects of parental and offspring treatments and their interaction. Differing letters above or below each group indicate significant pairwise differences in survival (Tukey’s HSD, *p* < 0.05).

### 3.2 Growth

Embryos reared under HypOA were 5.2% shorter at hatch (mixed-effect ANOVA, *p* < 0.001, Fig. 2E) and had yolk sacs that were 42% larger than control-reared embryos (mixed-effect ANOVA, *p* < 0.001, Fig. 2F). Parental treatment did not influence standard length of hatchlings (Fig. 2E). Yolk sacs were 14% larger in hatchlings from HypOA-treated parents when averaged across offspring treatments (mixed-effect ANOVA, *p* = 0.039), but pairwise comparisons did not find significant differences within offspring treatment (Fig. 2F). Normalized body depth at hatch was significantly smaller in offspring reared under HypOA (mixed-effect ANOVA, *p* = 0.045), but no significant differences were observed between individual treatment groups (Fig. 2G). On average, normalized eye width was significantly larger in hatchlings from HypOA-treated parents (mixed-effect ANOVA, *p* < 0.001, Fig. 2H). The parental × offspring treatment interactive effect was nearly significant (mixed-effect ANOVA, *p* = 0.051), as the effect of parental HypOA exposure was primarily driven by the relatively large eyes exhibited by the HC group (Fig. 2H).

At trial termination (19 dpf), larvae reared under HypOA were 12.9% shorter in length compared to larvae reared under control conditions (mixed-effect ANOVA, *p* < 0.001, Fig. 2I). This resulted in a 17.7% reduction in post-hatch growth rate in comparison to larvae reared in control conditions (mixed-effect ANOVA, *p* = 0.001, Fig. 2J). Additionally, the average growth rate of larvae from HypOA-treated parents was 10.8% lower relative to larval groups from control parents (mixed-effect ANOVA, *p* = 0.030, Fig. 2J). The interactive effect on growth rate was not significant, but the additive effect of parental and offspring HypOA exposure meant that HH larvae displayed the slowest overall larval growth, though not significantly different from CH larvae (Tukey’s HSD, *p* = 0.076, Fig. 2J). Normalized body depth was not statistically different between treatment groups (Fig. 2K). However, normalized eye width was significantly affected by both parental treatment (mixed-effect ANOVA, *p* = 0.042) and offspring treatments (mixed-effect ANOVA, *p* = 0.024, Fig. 2L). This effect was primarily driven by the HC group which exhibited larger eyes on average compared to all other groups (Fig. 2L).

### 3.3 Treatments effects on global gene expression

Analysis of RNAseq data with two-way PERMANOVA showed that global gene expression in silverside larvae was affected by both offspring (*p* < 0.001, R^2^ = 0.106) and parental treatment conditions (*p* = 0.008, R^2^ = 0.075, Fig. 3A). There was no interactive effect (Fig. 3A). Ordination using PCoA revealed distinct separation among the four treatment groups (Fig. 3A). Larvae reared under HypOA (CH and HH) are clustered away from the CC group, whereas larvae form the HC group replicates are positioned between the other clusters (Fig. 3A). In general, offspring from HypOA-treated parents clustered more closely than those from control parents (Fig. 3A). Intra-treatment transcript-wise variation in expression was greater within the CC and CH groups as compared to HC and HH larvae (Fig. 3B).

**Fig. 3.**
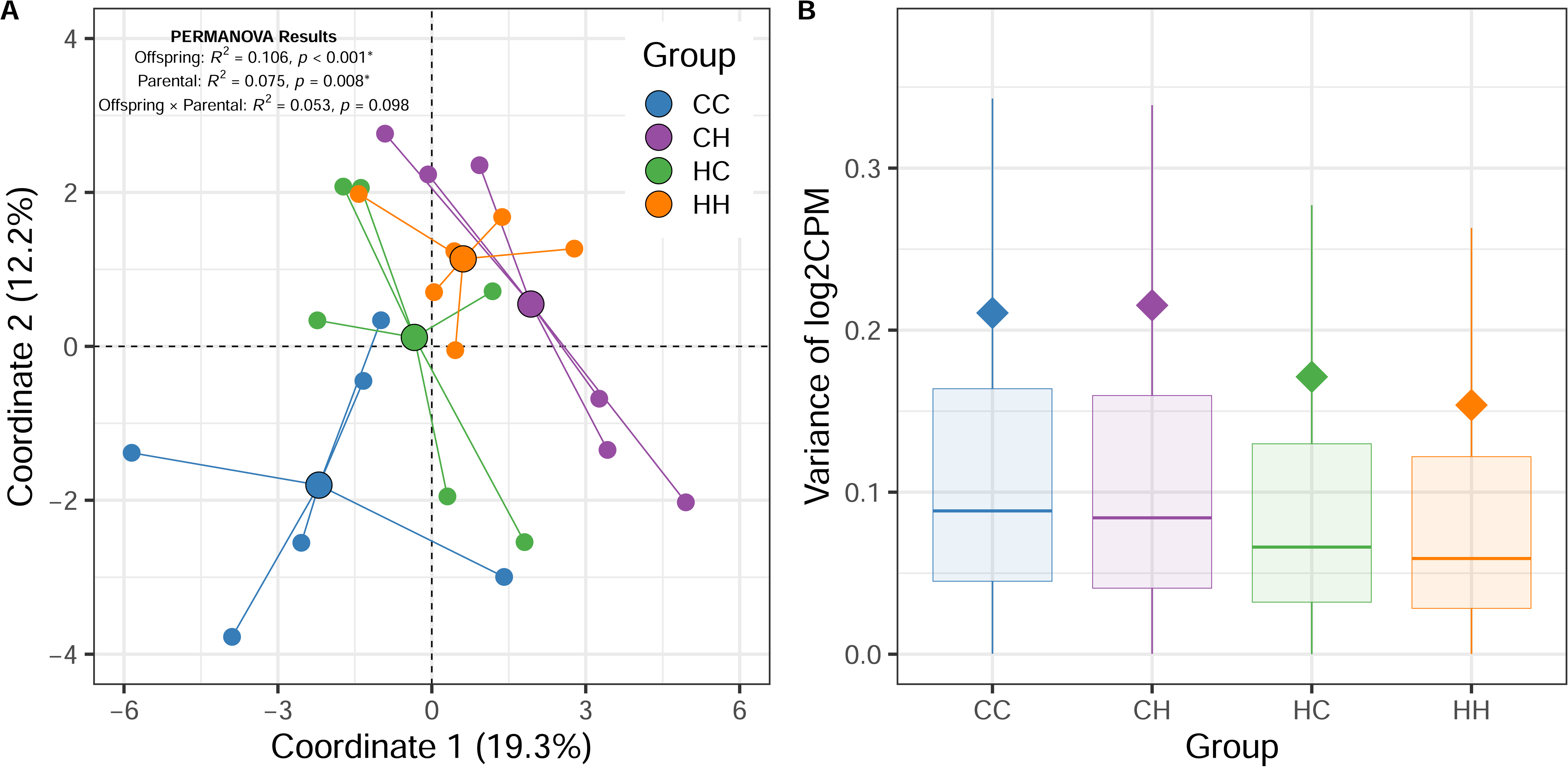
Treatment effects on global gene expression. (A) Results from principal coordinate analysis. Small colored circles represent the coordinates of individual samples (N =6), and they are connected to large colored circles, which represent the average coordinate position for each treatment group. The distances between points reflect the similarity in gene expression profiles among individual larvae and treatments. The results of a two-way PERMANOVA are presented in the top left of the panel, demonstrating the influence of both parental and offspring treatment conditions on the gene expression patterns observed. (B) Gene-wise variance of log_2_CPM across treatment groups. Colored boxes indicate the interquartile range, while the vertical lines extend to 1.5 times the interquartile range. The horizontal bar represents the median value for each treatment, and the colored diamond denotes the mean values. For clarity, individual variance values that fall outside the range are not shown in the plot.

### 3.4 Pairwise treatment tests of differential expression

Parental treatment had a strong effect on the number of DETs identified between control- and HypOA-reared larvae (Table 2, Fig. 4A). In offspring from control parents, HypOA exposure resulted in 1,606 DETs (CH vs. CC; Table 2). A similar number of DETs were found in HH group in comparison to CC larvae (1,520, Table 2). Although, more than half of these transcripts were distinct from those found in the CH vs. CC pairwise comparison (Fig. 4A-C). The were no DETs that were significantly upregulated in one group and significantly downregulated in the other. In contrast to larvae from control parents, larvae from HypOA parents exhibited just 4 DETs (HH vs HC; Table 2, Fig. 4A). Similarly, CH larvae exhibited just 58 DETs when compared to the HC group (Table 2, Fig4A). The two offspring groups reared under HypOA (HH vs CH) exhibited just three 3 DETs (Table 2) and there were no DETs found between the two offspring groups reared under control conditions (HCvs.CC, Table 2). Volcano plots for each pairwise comparison are presented in Fig S4. Supplementary Dataset 1 provides differential expression results for all transcripts and pairwise comparisons.

**Fig. 4.**
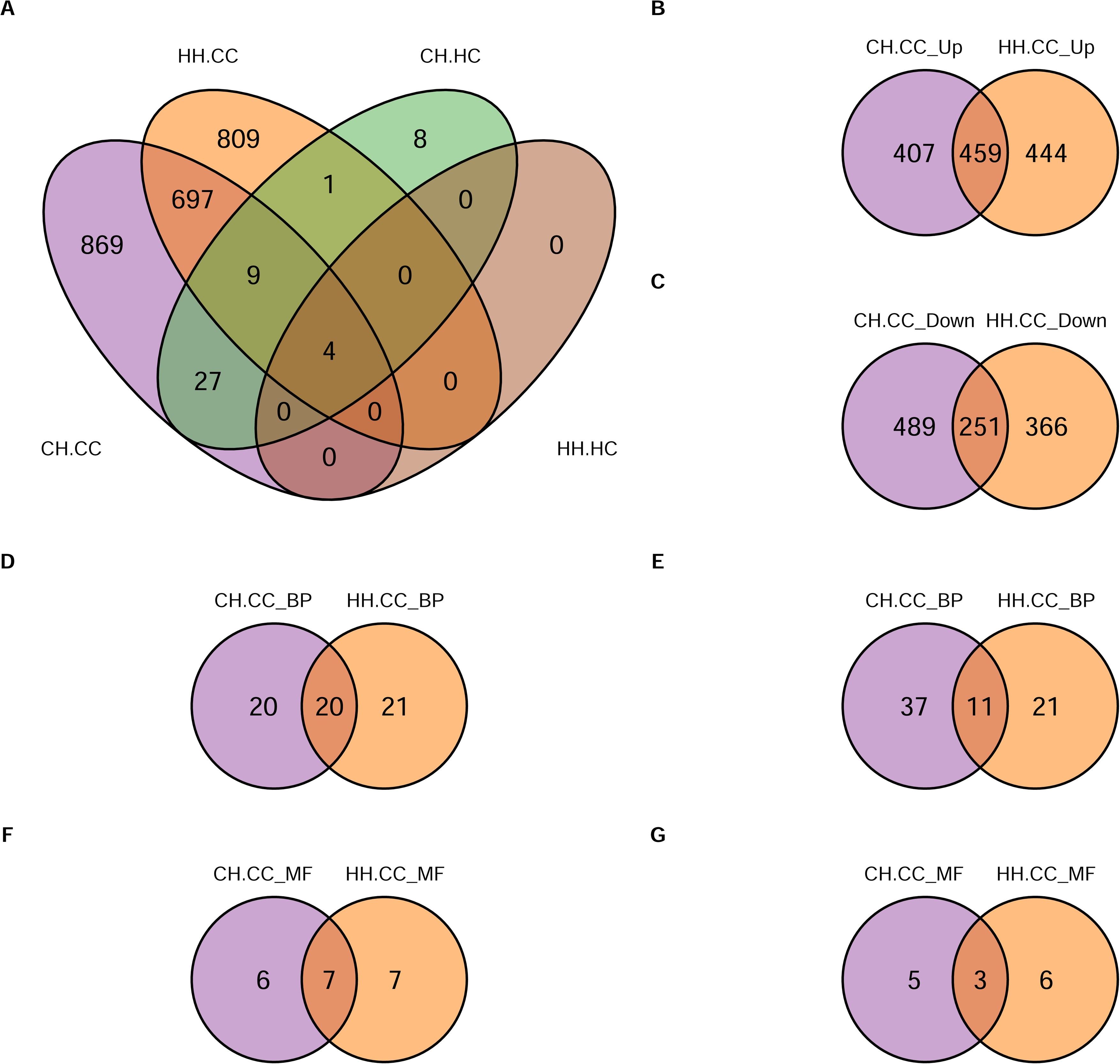
DET and GO Venn Diagrams. (A) Venn diagram showing shared and unique DETs from four pairwise treatment comparisons where significant DETs were detected (genewise negative binomial generalized linear models, FDR < 0.05). (B,C) Shared and unique DETs between the CHvs.CC and HHvs.CC pairwise comparisons for up- (B) and down-regulated DETs (C), respectively. (D-G) Venn diagrams showing the overlap of GO ‘biological process’ (BP) and ‘molecular function’ (MF) terms significantly enriched (*p* value < 0.05) in upregulated (D,F) and downregulated DETs (E,G) identified in larvae that were directly exposed to HypOA but from either control (CH) or HypOA-acclimated (HH) parental treatments.

**Table 2:**
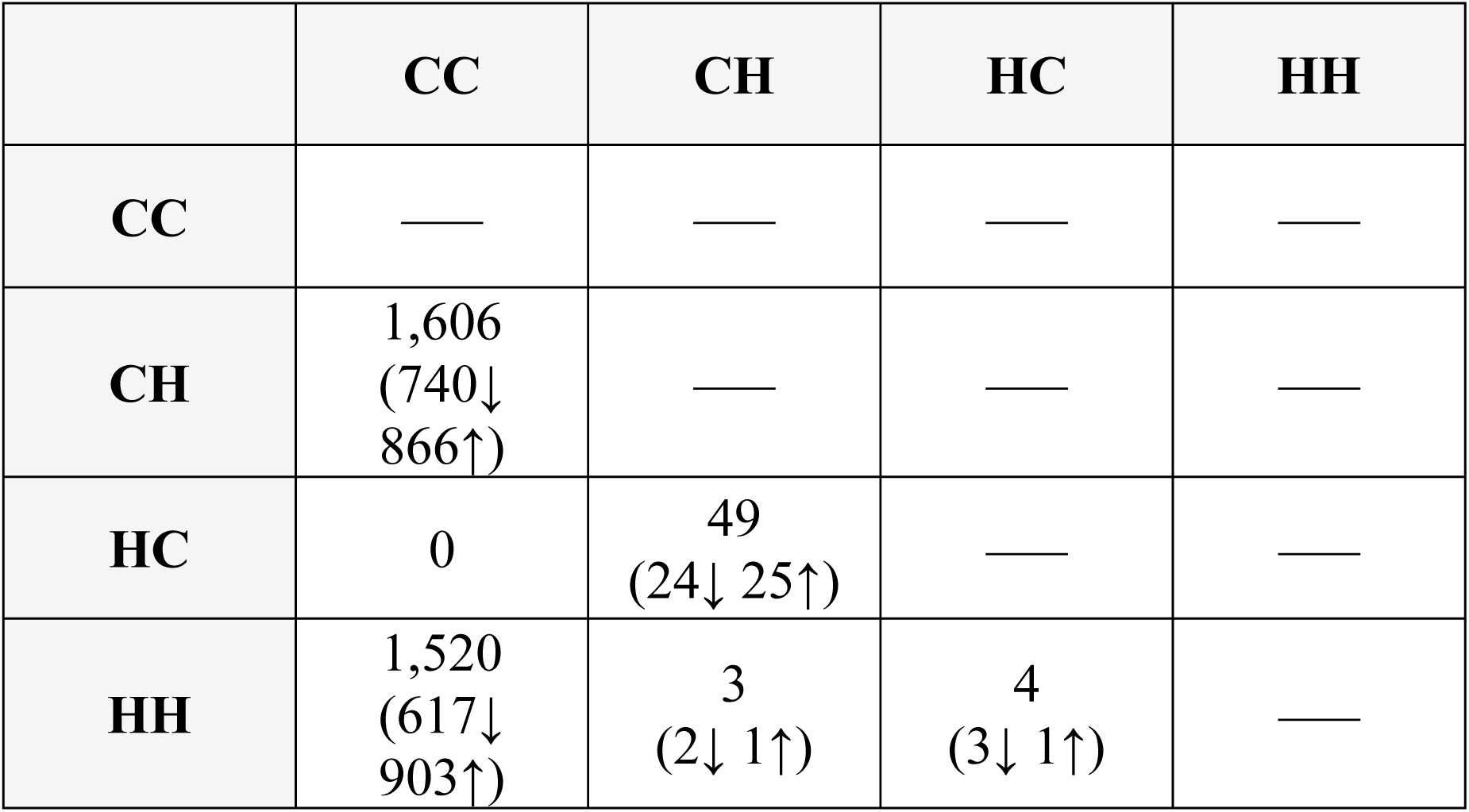
Matrix of the total number of differentially expressed transcripts between four treatment groups (N = 6 larval samples per treatment group). The number of up- and downregulated genes (in parentheses) refers to the row group compared to the column group.

|  | CC | CH | HC | HH |
| --- | --- | --- | --- | --- |
| CC | — | — | — | — |
| CH | 1,606<br>(740↓<br>866↑) | — | — | — |
| HC | 0 | 49<br>(24↓ 25↑) | — | — |
| HH | 1,520<br>(617↓<br>903↑) | 3<br>(2↓ 1↑) | 4<br>(3↓ 1↑) | — |

### 3.5 Functional enrichment of differentially expressed transcripts

GO enrichment analysis of up- and downregulated DETs from the CHvs.CC pairwise comparison found 88 GO: BP and 60 GO: MF terms. DETs from HHvs.CC pairwise comparison were enriched in 73 GO: BP and 25 GO:MF terms (see Supplementary Dataset 1 for detailed results). Many GO terms were unique to either CH or HH larvae (Fig. 4D-G). However, the most highly enriched terms were conserved between the two groups (Fig. 5). For example, upregulated DETs from both groups were enriched it GO: BP terms for neurotransmitter secretion, nervous system development, regulation of NMDA receptor activity, ionotropic glutamate receptor signaling pathway, and regulation of postsynaptic membrane potential (Fig. 5A,B). Top GO: MF terms in both groups included calcium channel regulator activity, calcium-dependent phospholipid binding, syntaxin-1 binding, and potassium channel activity (Fig. 5C,D). One difference was the enrichment of terms related to GABAergic and glutamatergic synaptic transmission in HH larvae, as several transcripts that regulate the expression of NMDA and GABA receptor subunits were significantly upregulated by HH larvae only (Fig. 5A,C). A similar pattern was found for downregulated DETs, in that the specific GO terms varied between groups (Fig. 4E,G), but the most highly enriched terms were shared between CH and HH larvae (Fig. 5). For example, the most enriched GO: BP terms for both groups included proteolysis, extracellular matrix organization, positive regulation of cell migration, and wound healing (Fig5A,B). Shared GO: MF terms included extracellular matrix constituent, serine-type endopeptidase activity, and heparin binding (Fig. 5C,D). Interestingly, there was 32% more GO terms enriched downregulated DETs for CH larvae compared to HH larvae (Fig. 4E,G), including unique terms related to immune response, such as phagocytosis and engulfment, response to bacterium, and viral entry into host (Dataset 1).

**Fig. 5:**
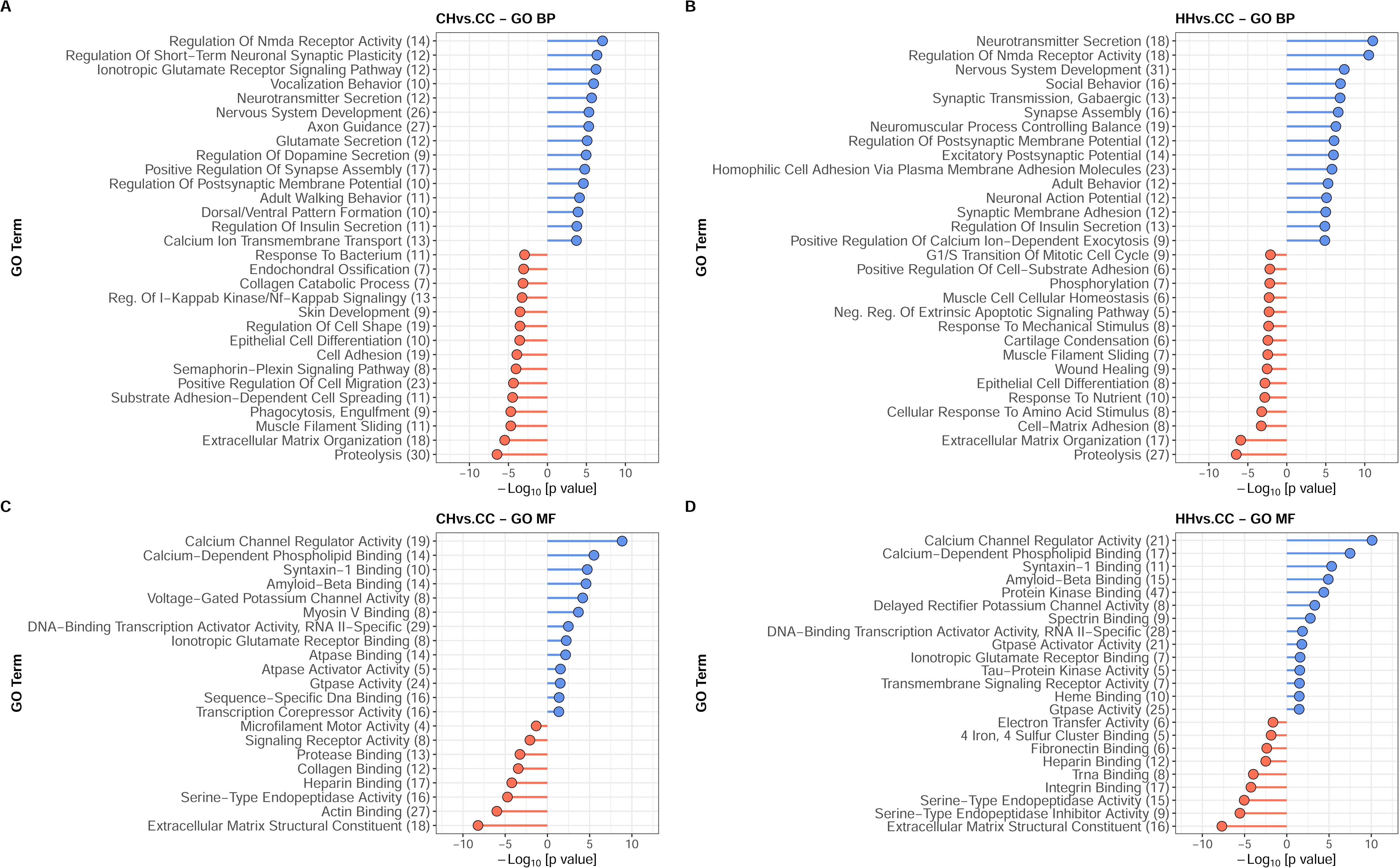
GO ‘biological process’ (BP) and ‘molecular function (MF) terms enriched in DETs expressed by CH and HH larvae. The length of the segment corresponds to the significance of the term (-log10[p-value] for up- (blue) and downregulated (red) DETs. Downregulated terms are shown with negative significance values for clarity. Values in parentheses next to term names indicate the number of transcripts contributing to the enrichment.

### 3.4 Effect of parental environment on larval transcriptional plasticity

While no DETs were detected between CC and HC larvae, we found that HC larvae exhibited a change in constitutive gene expression that matched the directional responses (i.e., up- or downregulated) of DETs expressed by larval groups that were directly exposed to HypOA, albeit to a lesser extent (Fig. 6A,B). This effect was highly consistent and held true for 95% of the 2,416 DETs that we analyzed for transcriptional frontloading (Fig. 6A,B). This indicates that parental exposure to HypOA alone had a sublet effect on the same molecular functions as direct offspring exposure to HypOA.

**Fig. 6:**
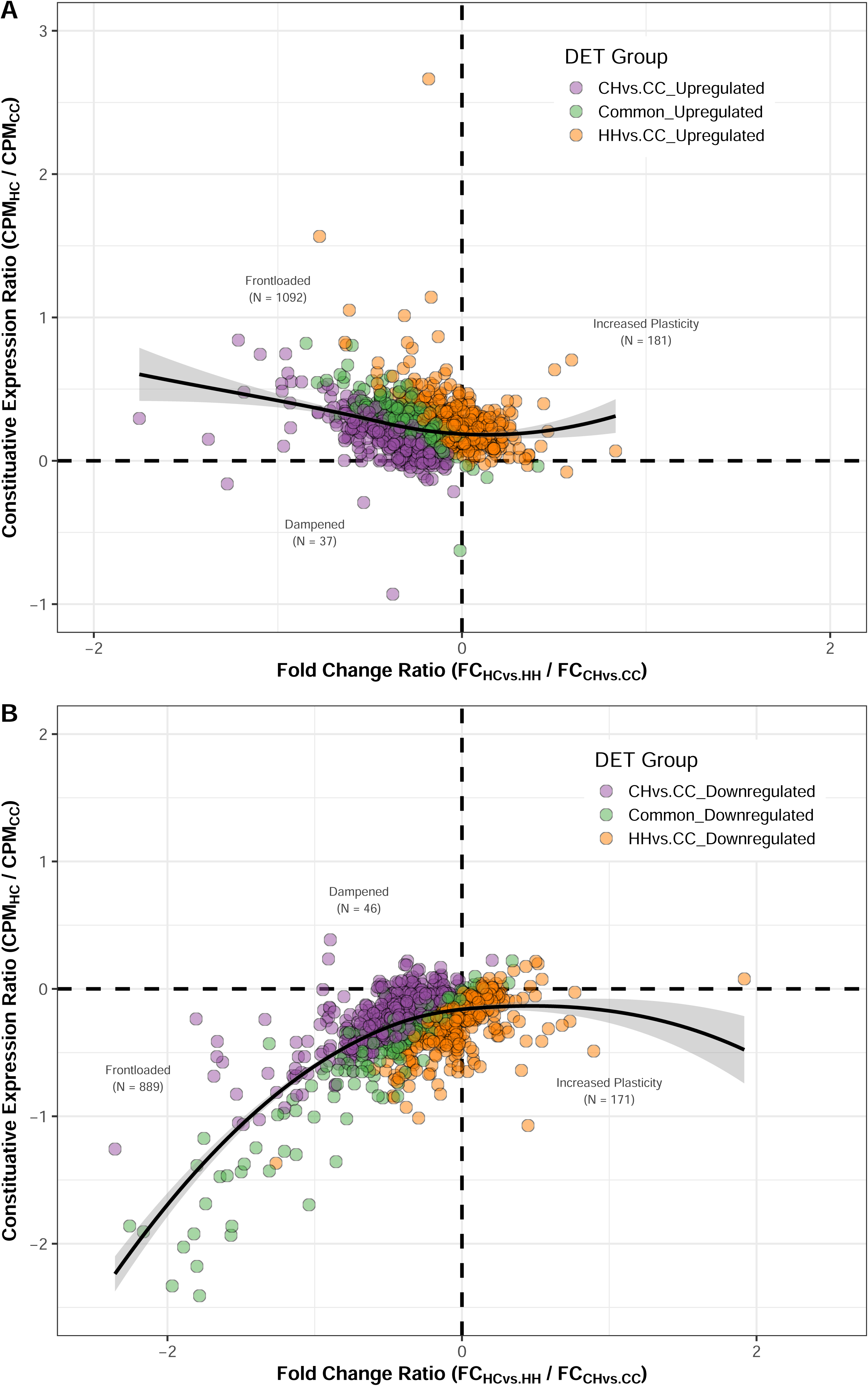
Patterns of transcriptional frontloading. Scatterplots comparing the fold change ratio (x-axis) to the constitutive expression ratio (y-axis) across 2,416 differentially expressed upregulated (A) and downregulated (B) transcripts. Individual transcripts were binned into three categories: frontloaded (shaded quadrant), dampened, or increased plasticity. Colored circles indicate whether genes were commonly different expressed in both CH and HH larvae or were uniquely expressed in either CH or HH larvae. Scatter plots are supplemented with a LOESS curve (solid black line ± 95% CI).

#### ‘Frontloaded’ Transcripts

The majority of these DETs were categorized as ‘frontloaded’ in larvae from HypOA-treated parents (82% of total DETs, Fig. 6A,B). The expression of frontloaded DETs was significantly affected by a parental treatment × offspring treatment interactive effect (ANOVA; *p* < 0.001) and results from pairwise comparison tests supported the frontloading criteria (Fig. 7A,D). Compared to CC larvae, mean expression in HC larvae was significantly greater for upregulated DETs (Tukey’s HSD; p < 0.001, Fig. 7A) and significantly lower for downregulated DETs (Tukey’s HSD; p < 0.001, Fig. 7D). As a result, the change in gene expression between groups reared under control and HypOA conditions was reduced by over three-fold in larvae from HypOA-treated parents compared to larvae from control parents (Fig7A,D). Frontloaded DETs were enriched in the same top GO: BP and MF terms found for GO enrichment of CH and HH larvae (Fig. S5).

**Fig. 7:**
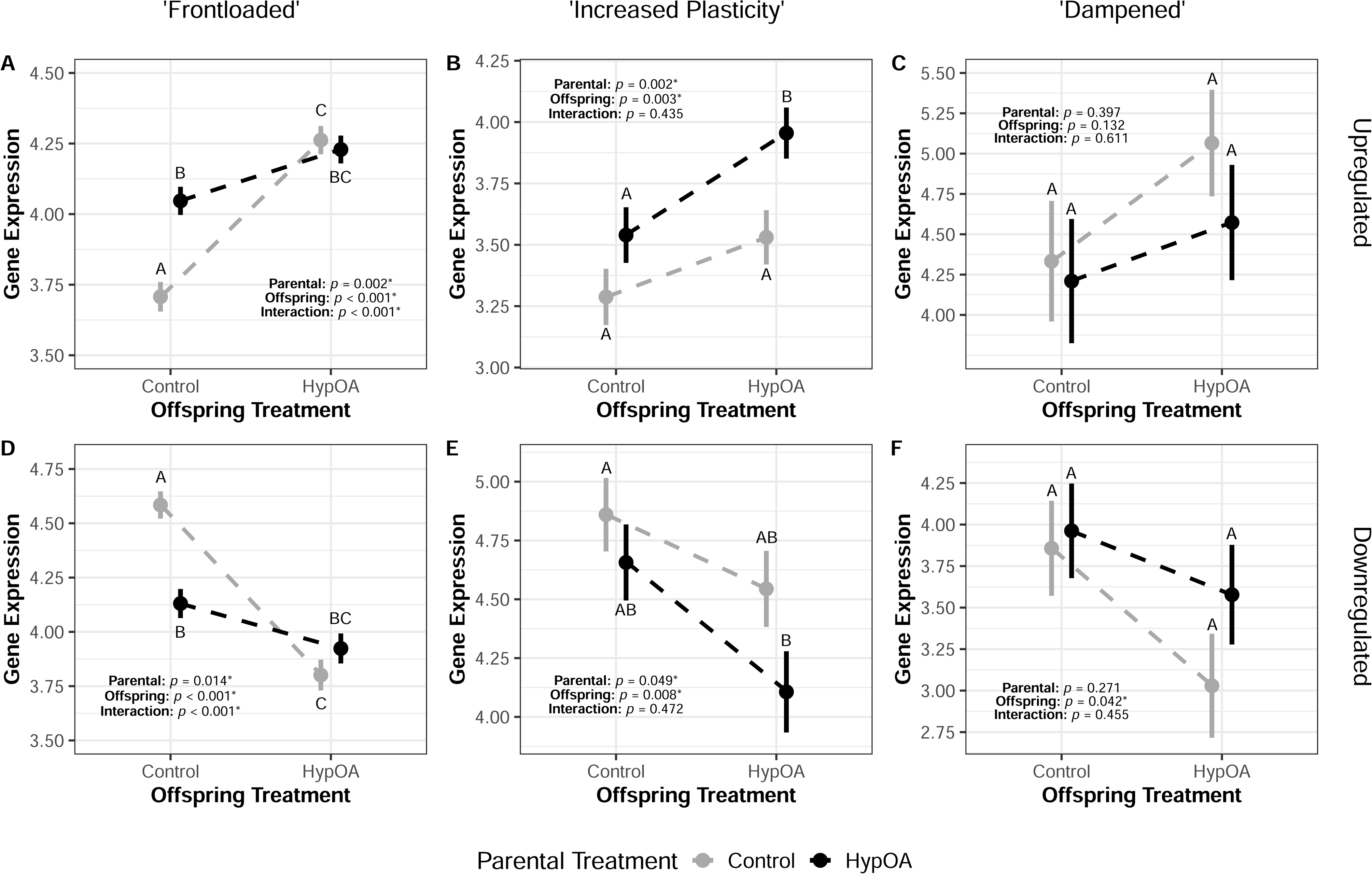
Gene expression reaction norms for HypOA exposure by parental treatment. Larval gene expression (normalized and variance stabilized log_2_CPM) of ‘frontloaded’ (A,D), ‘increased plasticity’ (B,E), and ‘dampened’ genes (C,F) genes that were up- or downregulated in larvae that were reared in the HypOA treatment. White and black circles represent mean expression by treatment group (vertical lines denote ±1 s.e.). Hashed lines represent the larval reaction norms by parental treatment. Significance levels (*p* values) are presented from two-way ANOVAs testing for treatments effects on gene expression of within different categories. Differing letters indicate significant pairwise differences (Tukey’s HSD, *p* < 0.05) between treatment groups.

#### ‘Increased Plasticity’ Transcripts

We found that 14.6% of DETs fit the criteria for ‘increased plasticity’ (Fig. 6A,B). For these DETS, larvae from HypOA-treated adults showed a larger change in expression between offspring treatments compared to the response of larvae from control-treated parents (Fig. 6A,B). As a result, HH larvae exhibited the highest mean expression among upregulated DETs (Fig. 7B) and the lowest mean expression among downregulated DETs (Fig. 7E). Nearly all ‘increased plasticity’ DETs (94.8%) were identified within the HHvs.CC pairwise comparison only (Fig. 6A,B), suggesting that the increased responsiveness of these genes was specifically linked to parental exposure to HypOA. There were 19 GO: BP and 5 GO:MF terms enriched in upregulated ‘increased plasticity’ DETs (Dataset 1, Fig. S5), including high enriched terms for brain development and excitatory post synaptic potential. There were 15 GO: BP and 4 GO:MF terms enriched in downregulated ‘increased plasticity’ DETs (Dataset 1, Fig. S5). Top enriched terms included cell redox homeostasis, skeletal muscle cell differentiation, and chromatin organization.

#### ‘Dampened’ Transcripts

A small number of DETs were categorized as ‘dampened’ (3.4% of total DETs, Fig. 6A,B). In general, larvae from HypOA-treated parents expressed upregulated DETs at lower levels and downregulated DETs at higher levels, although no significant differences were found (Fig. 7C,F). Dampened transcripts that were upregulated under HypOA were enriched in 6 GO: BP and 3 GO: MF terms enriched upregulated DETs (Dataset 1, Fig. S5). Downregulated dampened DETs were enriched in 9 GO: BP and 5 GO: MF terms (Dataset 1, Fig. S5).

## 4. Discussion

### 4.1 Cross-generational plasticity to HypOA mediated through transcriptional frontloading

The increasing severity of coastal hypoxia and acidification (HypOA) threatens fish populations that depend on productive nearshore environments for reproduction (Gobler and Baumann, 2016). Fish have evolved various behavioral and physiological adaptations to low DO and elevated *p*CO_2_ (Esbaugh, 2018; Richards, 2009). However, few studies have examined the underlying changes in gene expression that facilitate developmental responses to this combined stressor. Furthermore, cross-generational plasticity may play an important role in priming adaptive responses in offspring. Anticipatory cross-generational plasticity is thought to evolve in populations inhabiting environments with regular environmental fluctuations and where parental conditions are generally predictive of the offspring environment (Bonduriansky, 2021). Given that HypOA conditions are common in temperate coastal systems (Baumann and Smith, 2017), if a parent is exposed to HypOA before spawning, it heightens the probability that offspring will encounter these conditions during development.

In this study, we investigated the potential for anticipatory cross-generational plasticity in the Atlantic silverside, a coastal forage fish that utilizes temperate nearshore systems as spawning and nursery grounds. We found that offspring from control parents that were reared under HypOA exhibited typical declines in survival and developmental rates. By contrast, the survival of embryos fertilized by HypOA-treated parents was not affected by the HypOA treatment. Though, the overall survival and post-hatch growth rates of larvae from HypOA-treated parents was, on average, lower than larvae from control parents. Using RNAseq, we found that direct exposure to HypOA induced extensive changes in larval gene expression. We identified several impacted molecular pathways, including those regulated by the hypoxia-induced factor (Hif) signaling pathway and other well-described regulatory pathways. The expression of mRNAs encoding Hif proteins themselves were not significantly affected by treatment conditions (Fig. 8A). This is not surprising given the long-term nature of our HypOA treatment and the oxygen-dependent post-translational regulation of Hif activation (Semenza, 2000). Instead, we found several affected processes downstream of Hif that control nervous system development, axonal pathfinding, neurotransmitter regulation, and extracellular matrix homeostasis (Bonkowsky and Son, 2018; Mandic et al., 2021; Semenza, 2000).

**Fig. 8:**
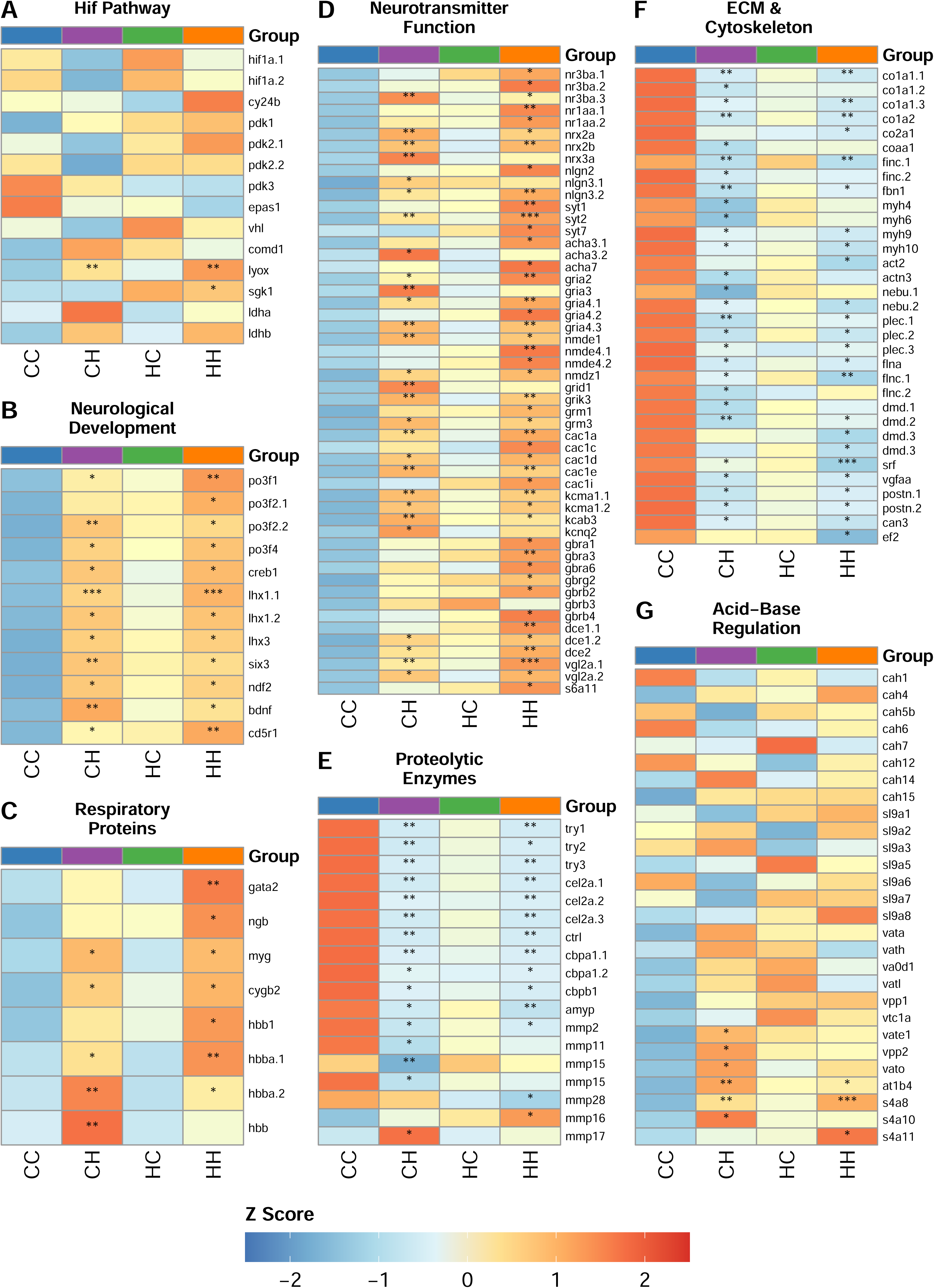
Heatmaps of gene expression for selected cellular pathways. Heatmaps showing gene expression (log_2_CPM normalized Z-score) across four treatment groups within relevant transcriptional pathways. Rows show individual transcripts that are labeled by gene symbol on the right (.1, .2, .3 indicate multiple isoforms for a single gene symbol). Stars inside the cell indicate that expression in that group was significantly different than CC larvae, the overall control group (* FDR < 0.05, ** FDR < 0.01, *** FDR < 0.001). (A) Hif Pathway; (B) Neurological Development; (C) Neurotransmitter Function; (D) Extracellular matrix (ECM) and Cytoskeleton; (E) Proteolytic Enzymes; (F) Respiratory Proteins; (G) Acid-Base Regulation.

Furthermore, larvae from HypOA-treated parents that were reared under control conditions exhibited a pattern of constitutive gene expression that was similar to larvae that were directly exposed to HypOA. While these changes in expression were subtle and did not reach statistical significance, they were consistent across hundreds of genes that were significantly differentially expressed by direct HypOA exposure. This pattern broadly aligns with the framework of transcriptional frontloading, a type of acclimatory gene expression commonly observed in marine animals that inhabit variable environments and are repeatedly exposed to periods of abiotic stress (Barshis et al., 2013; Collins et al., 2021; Gurr et al., 2022). Our results demonstrate that a similar transcriptional response can arise in offspring when their parents are pre-exposed to a priming stressor. We interpret this pattern as a pre-activation of the molecular stress response to HypOA, which may enhance the capability of offspring to adequately sense and respond to direct exposures (Hackerott et al., 2021; Sopinka et al., 2017). For example, we found that larvae from HypOA-treated parents showed increased expression of protein-lysine 6-oxidase (Fig.10A), which is an important marker to Hif activation (Lin et al., 2020). Furthermore, serum and glucocorticoid-regulated kinase 1 was upregulated by more than 12-fold in larvae from HypOA-treated parents (Fig. 8A), which is activated by cortisol and is a key factor that coordinates various cellular stress response mechanisms (Notch et al., 2011).

### 4.2 Transcriptional Effects Associated with Neurological Development

The developing central nervous system is highly vulnerable to hypoxia, as insufficient oxygen levels impair critical developmental processes related to neurogenesis, axon pathfinding, and synapse formation (Bonkowsky and Son, 2018). Reduced aerobic energy production and a loss of ion regulatory capacity leads to increased rates of neuronal cell death via apoptosis of necrosis (Banasiak et al., 2000). For example, zebrafish larvae exposed to a DO level of 10% a.s. displayed decreased neural proliferation in the forebrain and a downregulation of genes associated with the development of neurons, glia, and oligodendrocytes (Mikloska et al., 2022). However, in this study we exposed silverside offspring to a more moderate hypoxic treatment (40% a.s.) and observed upregulation of genes associated with nervous system development, neurotransmitter activity, and regulation of neuron membrane potential through increased expression of calcium ion channels and voltage-gated potassium channels (Fig. 8B,D).

Moreover, these pathways were broadly frontloaded in larvae from HypOA-treated parents, involving several transcription factors that coordinate nervous system development (Fig. 8B). Frontloaded transcription factors include, but are not limited to, POU-domain class 3 transcription factors (*pou3f1, pou3f2, pou3f3b*, and *pou3f4*) (Latchman, 1999), cyclic AMP-responsive element-binding protein 1 (Lee et al., 2018), LIM/homeobox protein 3 (Hobert and Westphal, 2000), homeobox protein six3 (Chao et al., 2010), neurogenic differentiation factor 2 (Olson et al., 2001), brain-derived neurotrophic factor protein (Anand and Mondal, 2020), and cyclin-dependent kinase 5 activator 1 (also known as *p35*), which has anti-apoptotic effects and has been shown to regulate neurogenesis and development of the retina in zebrafish (Leung et al., 2008). The upregulation of these genes suggests a protective response within the central nervous system against further insults. Like other components on the central nervous system, the development of the retina is sensitive to hypoxia, and even modest reductions in available oxygen can impact visual performance (McCormick and Levin, 2017). Hence, smaller eyes may be an expected phenotypic response of larvae reared in oxygen limiting conditions. However, we observed that silverside larvae that were directly exposed to HypOA maintained a normal eye size relative to their body length. Additionally, newly hatched larvae from HypOA-treated parents showed larger eyes than those from control parents, especially in larvae reared under control conditions, which developed larger eyes than all other treatment groups. Our results suggest that silverside offspring can alter the expression of genes involved in neurological and sensory development to support eye growth under HypOA conditions. Furthermore, the frontloading of these genes in offspring from HypOA-treated parents led to a measurable overgrowth in eye size when the HypOA stressor was removed. Silverside larvae are primarily visual particulate feeders (Gilmurray and Daborn, 1981) and rely on the rapid development of visual systems to locate prey and avoid predation. These findings necessitate further research to evaluate the functional impacts on vision resulting from cross-generational exposure to HypOA.

### 4.3 Within- and Cross-generational Effects of HypOA on Neurotransmitter Function

Larvae reared under HypOA exhibited extensive upregulation of genes associated with synapse formation and neurotransmitter regulation. This involved increased expression of several neurexin and neuroligin isoforms (Fig. 8D), which are proteins that form the trans-synaptic complexes between pre- and postsynaptic neurons and facilitate synapse formation and neurotransmitter release (Bottos et al., 2011). We also found an upregulation of several synaptotagmin isoforms (Fig. 8D), which are trafficking proteins located in the presynaptic membrane that regulate rapid neurotransmitter release (Chapman, 2008). Furthermore, exposed larvae showed increased expression of genes involved in excitatory neurotransmitter signaling, including acetylcholine receptors, and other ionotropic and metabotropic glutamate receptors (Fig. 8D). This pattern was accompanied by the upregulation of mRNAs encoding voltage-dependent calcium ion channels and voltage-gated potassium ion rectifier channels (Fig. 8D). Excitatory neurotransmission is vital for neuronal development and memory formation; however, secreted glutamate becomes excitotoxic under hypoxic conditions (Nilsson, 1995). Anoxia-tolerant species like Crucian carp (*Carassius Carassius*) and goldfish (*C. auratus*) can survive extreme low oxygen, in part, through depression of neurological activity. This is achieved through increased production of gamma-aminobutyric acid (GABA), the major inhibitory neurotransmitter in fish brain, which alleviates neuronal excitotoxicity by inhibiting neurotransmission and promoting cell survival through anti-apoptotic and anti-oxidative effects (Nilsson, 1995). Interestingly, genes involved in GABA neurotransmission were broadly frontloaded and expressed a greater level in larvae from HypOA-treated parents, including genes encoding GABA_A_ receptors, GABA_b_ receptors, and proteins involved in GABA synthesis and transport (Fig. 8D).

Whether transcriptional frontloading and increased expression of these genes contributed to the different phenotypic responses of offspring from HypOA-treated parents is an important area for further investigation. Especially considering that exposure to elevated *p*CO_2_ can alter the function of GABAergic neurotransmission in fish. Under acidification, acid-base regulatory adjustments alter ion gradients across neuronal membranes and cause a reversal of function in some GABA_A_ receptors, resulting in excitatory instead of inhibitory signaling (Heuer et al., 2019; Nilsson et al., 2012). This effect has been linked to behavioral dysfunction in some tested fish species (Schunter et al., 2019). Similar effects to olfaction and other aspects of sensory perception have been documented, indicating that exposure to elevated *p*CO_2_ may have broader impacts on neurological function (Cohen-Rengifo et al., 2022; Heuer et al., 2019). Interestingly, parental exposure to elevated *p*CO_2_ has been shown to offset behavioral effects in offspring, suggesting that transcriptional flexibility can restore normal GABA function (Schunter et al., 2016). Our findings offer further support that cross-generational exposure to HypOA can alter gene expression patterns around neurotransmitter regulation and function in fish early life stages.

### 4.4 Cross-generation Exposure to HypOA Amplified Metabolic Depression

Metabolic depression is a primary strategy employed by fish to survive periods of low DO (Richards, 2009). For example, the anoxia-tolerant embryos of the annual killifish (*Austrofundulus limnaeus*) reduce metabolic rate in an oxygen dependent manner and do not increase ATP production through anaerobic metabolism (Anderson and Podrabsky, 2014). In this study, Atlantic silverside offspring responded to HypOA in a similar manner by exhibiting delayed development, reduced size at hatch, and slower rates of post-hatch larval growth. Furthermore, transcripts encoding proteins required for anaerobic energy production (e.g., lactate dehydrogenases) were not significantly affected by parental or offspring exposure to HypOA (Fig. 8A). Instead, we found a general downregulation of genes involved in somatic growth and tissue development. Specific mRNAs included major components of the extracellular matrix, such as collagens, fibronectin, and fibrillin, as well as cytoskeleton components and other proteins involved in muscle development and contraction, including myosin heavy chains, actins, nebulins, plectins, filamens, and dystrophins (Fig. 8F). This pattern was accompanied with a downregulation of growth factors and proteins implicated in tissue remodeling and muscle development, including serum response factor (Miano et al., 2007), periostin (Kudo et al., 2004), and calpain-3 (Prykhozhij et al., 2023) (Fig. 8F). In part, this transcriptional pattern may reflect that larvae reared under HypOA were smaller and less developed than control larvae at the time of sampling. However, the fact the suppressive pattern of these genes was frontloaded in larvae from HypOA-treated parents suggests an adaptive response to hypoxia. Downregulation of these transcripts allows for metabolic savings by reducing the cellular machinery needed to support muscle development and movement, as well as a general reduction to the synthesis of these highly abundant proteins (Richards, 2009). Similar adaptive responses to hypoxia related to muscle contraction proteins, extracellular matrix components, and cytoskeletal proteins have been observed in long-jawed mudsucker (*Gillichthys mirabilis*) and zebrafish (Gracey et al., 2001; Ton et al., 2003).

### 4.5 Frontloaded Expression of Respiratory Proteins

Fish exposed to hypoxia typically upregulate the expression globin proteins that bind oxygen and transport it to demanding tissues (Richards, 2009). We found that mRNAs encoding several globin proteins were strongly upregulated in larvae reared under HypOA (Fig. 8C). Several of these genes were positively frontloaded in larvae from HypOA-treated adults, including myoglobin, cytoglobin-2, and a transcript annotated for hemoglobin subunit beta-1, which was the most abundantly expressed hemoglobin mRNA by over 40-fold (Fig. 8C). We also found that neuroglobin expression was strongly frontloaded in larvae from HypOA-treated parents (Fig. 8C). This globin protein is localized in neurons and protects the brain from ischemic injury by rapidly detecting hypoxic conditions and enhancing oxygen delivery (Fuchs et al., 2004). Interestingly, the parental effects on the expression of respiratory proteins were accompanied by a positive frontloading of the *gata2* transcription factor in larvae from HypOA-treated parents (Fig. 8C). *gata2* regulates the differentiation and maintenance of hematopoietic stem cells in fish (Gioacchino et al., 2021). A similar cross-generational response was observed in zebrafish after paternal treatment to hypoxia, leading to heightened offspring expression of hemoglobin proteins that was associated with enhanced hypoxic tolerance (Ragsdale et al., 2022). Our findings corroborate the finding that parental exposures to hypoxia can alter gene expression patterns around the regulation of erythropoiesis and hemoglobin synthesis, potentially increasing offspring survival to hypoxic conditions.

### 4.6 Transcriptional responses to elevated *p*CO_2_

Transcriptional responses in fish to elevated *p*CO_2_ are less understood compared to the effects of hypoxia and other stressors like warming (Baumann, 2019). The combined HypOA treatment employed in this study prevents us from isolating the individual transcriptional effects of elevated *p*CO_2_. Though, we can still analyze the expression genes that are critical for effective acid-base regulation. For example, carbonic anhydrases facilitate CO_2_ excretion, ionic regulation, and acid–base balance by catalyzing the hydration of CO_2_ into H^+^ and HCO_32-_ for export out of the body (Gilmour and Perry, 2009). However, we found that mRNA levels of carbonic anhydrase isoforms with catalytic activity were not significantly affected by parental or offspring treatments (Fig. 8G). Larval fish primarily regulate acid/base balance through cutaneous ionocytes, which express high levels of membrane-based ion pumps and co-transporter proteins (Dahlke et al., 2020). Again, we found many of these proteins were unaffected by treatment conditions, including sodium/hydrogen exchangers and vacuolar proton-transporting V-type ATPases (Fig. 8G). Furthermore, sodium/potassium ATPases were mostly downregulated under HypOA (Fig. 8G). A few vacuolar proton-transporting V-type ATPase subunits were slightly but significantly upregulated in CH larvae only, though these transcripts were positively frontloaded in larvae form HypOA-treated parents (Fig. 8G). We also found a slight upregulation of several chloride/sodium/bicarbonate exchangers (Fig. 8G), that have putative roles in acid/base regulation (Bayaa et al., 2009).

These results indicate that Atlantic silverside larvae, like many other tested species, exhibit adequate acid/base regulatory capacity under basal levels of gene expression to successfully buffer against environmentally relevant levels of acidification (Esbaugh, 2018; Kwan et al., 2021). On the other hand, ion pumping is an energy-demanding process, and the need for increased acid/base regulatory capacity to counter acidification conflicts with the requirement for metabolic depression under hypoxia (Perry and Gilmour, 2006; Richards, 2009). In this scenario, fish larvae may suffer a heightened risk of acidosis when simultaneously exposed to low DO, an effect that may contribute to negative interactive effects observed in Atlantic silverside offspring (DePasquale et al., 2015; Miller et al., 2016; Schwemmer et al., 2020). Targeted experimental designs are required to clarify how hypoxia alters the transcriptional response to seawater acidification in fish early life stages.

### 4.7 Conclusion

We demonstrated that Atlantic silverside offspring respond to HypOA through extensive upregulation of genes associated neurological development, synaptic plasticity, neurotransmitter function, and ion regulation in the central nervous system. The energetic costs associated with protecting the development of the central nervous system may be balanced by a reduction in the expression of genes that promote somatic growth and muscle development. Furthermore, we showed that parental exposure to HypOA prior to spawning directly influences how offspring respond to this common abiotic stressor. We observed a pattern of cross-generational transcriptional frontloading, wherein genes responsible for mediating the molecular response to HypOA showed a change in constitutive expression that matched the pattern observed in larvae that were directly exposed to HypOA. Transcriptional frontloading may have enhanced the capacity of offspring to effectively respond to HypOA conditions, as evidenced by increased embryonic survival.

Importantly, our findings illustrate the potential of cross-generational plasticity in facilitating adaptive responses to emerging anthropogenic stressors, potentially enabling populations to reproduce and persist in impacted environments in lieu of rapid genetic adaptation (Munday, 2014). There are several potential mechanisms mediating cross-generational plasticity, including many maternal effects and epigenetic factors that can alter offspring gene expression (Adrian-Kalchhauser et al., 2020; Green, 2008). It is probable that these individual factors act simultaneously through complex synergistic or antagonistic interactions that can result in both beneficial and deleterious effects in offspring. For example, we found that larvae from HypOA-treated parents that were reared under control conditions showed lower survival and post-hatch growth compared to conspecifics for control parents. This suggest that parental exposure to HypOA involved the transfer of stress response components that diminish offspring fitness (i.e., a condition transfer) (Bonduriansky, 2021). Maternal stress is known to increase the level of maternal cortisol transferred to embryos, resulting in significant changes in neurological development and behavior (Best et al., 2017). Increased loading of maternal hormones like cortisol may prepare offspring for stressful conditions but can also promote maladaptive phenotypes in offspring (Sopinka et al., 2017). Subsequent research should prioritize investigating the precise non-genetic mechanisms mediating cross-generational plasticity, particularly during embryonic development. Research is also needed to characterize the persistence of transcriptional frontloading in response to parental or developmental stress and the long-term implications for organismal fitness within and across generations.

## List of Symbols and Abbreviations

a.s.: air saturation
*A*_T_: total alkalinity
ANOVA: analysis of variance
BP: biological process term (Gene Ontology)
CC: Control-treated parents and control-reared offspring
CH: Control-treated parents and HypOA-reared offspring
CPM: counts per million
ESL: Environmental Systems Laboratory
DET: differentially expressed transcript
DIC: dissolved inorganic carbon
DO: dissolved oxygen
dpf: days post-fertilization
dph: days post-hatch
FC: fold change
FDR: false discovery rate
GABA: gamma-aminobutyric acid
GO: Gene Ontology
HC: HypOA-treated parents and control-reared offspring
HH: HypOA-treated parents and HypOA-reared offspring
Hif: hypoxia induced factor
HypOA: concurrent hypoxia and acidification
IVF: *in vitro* fertilization
MF: molecular function term (Gene Ontology)
*p*CO_2_: partial pressure of CO_2_
pH_T_: pH on the total scale
WHOI: Woods Hole Oceanographic Institution

## Acknowledgements

We are grateful to Dr. Adam Subhas and Matt Hayden for conducting carbon chemistry measurements. We thank Solomon Chen for assistance in programing the automated HypOA system.

## Competing Interests

No competing interests declared.

## Funding

Financial support for this study and C.S.M was provided by a National Science Foundation Postdoctoral Fellowship in Oceanography (Award Number: 2126533). Support for A.M. was provided by the Woods Hole Partnership in Education Program. N.A. and M.L. were not funded by a specific funding source for this project.

## Data Availability Statement

Raw sequence reads and HTseq-count output files are deposited in NCBI GEO Database (accession number: GSE245490). Use this token for reviewer access: cholwsmcfhovhkv. Datasets for survival (http://lod.bco-dmo.org/id/dataset/925081) and growth (http://lod.bco-dmo.org/id/dataset/925042) are deposited in a public BCO-DMO repository.

## Notes

### Competing Interest Statement

The authors have declared no competing interest.

